# Programming primary human T cells to deliver gene editing machinery

**DOI:** 10.64898/2026.09.30.753584

**Authors:** James Meixiong, Oleksandr Zginnyk, Lauren Chow, Brian R. Shy, Matthew H. Spitzer

## Abstract

Clinical applications of gene editors including CRISPR-Cas9 are limited by the cell-type specificity and tissue penetrance of current *in vivo* delivery options. Cell-mediated gene editor delivery could overcome these obstacles. To date, systems utilizing immortalized cell lines have demonstrated gene editor delivery *in vitro*. Because T cells traffic throughout the organism and engage in intercellular communication with individual target cells, we wondered whether they could be engineered to deliver gene editing machinery. Here, we demonstrate that primary human T cells can produce retroviral-based enveloped delivery vehicles (EDVs) packaging functional Cas9 ribonucleoproteins. When paired with fusogenic VSV-G, T cell-generated Cas9-EDVs edited a diverse array of target cells. We developed a miniaturized Moloney murine leukemia virus-based system that proved superior to existing architectures in this context and optimized both lentiviral and site-specific knock-in approaches for T cell engineering. These Enveloped Particle Information Courier-(EPIC-) T cells offer key advantages as delivery systems.

## INTRODUCTION

Since their discovery and characterization, CRISPR-Cas9 and other RNA-guided nuclease systems have rapidly transformed scientific research and spurred global biotechnological innovation for gene editing applications. *In vitro*, these technologies enable installation of bespoke edits at genome scale. Despite these capabilities, *in vivo* delivery remains a significant barrier for clinical adoption^1,2^. FDA-approved *in vivo* delivery vehicles for gene editors primarily include lipid nanoparticles and viral vectors, including adeno-associated virus (AAV) and lentivirus, each with unique strengths and weaknesses. Lipid nanoparticles (LNPs) can deliver nucleases as mRNA or ribonucleoproteins (RNPs), both of which are less genotoxic than DNA-encoded nucleases. Despite proven safety and scalability, their trafficking capabilities are generally limited to select organs, and off-target cell populations such as endothelial cells act as large sinks for the administered dose^3-6^. In the case of viral vectors, tropism can be engineered via capsid (AAV) or membrane proteins (lentivirus) to direct cell-type specific recognition and fusion^7,8^, but viral options suffer from oncogenic risk from transgene insertion^2^, genotoxicity from DNA-encoded nucleases, and immunogenicity, which hinders repeat dosing and risks rejection of edited target cells.

An ideal delivery system that merges the strengths of current methods would be an intravenous agent that delivers a transient gene editor with cell-type specificity. Cell-based delivery represents one promising solution that meets these criteria. Proposed cell-based approaches include cells shuttling oncolytic virus that encodes gene editors^9^, transferring gene editors to target cells via trogocytosis of membrane associated proteins^10^, and producing retroviral-based protein nanocages^11^, exosomes/extracellular vesicles^12,13^, or enveloped delivery vehicles (EDVs, also known as viral like particles, or VLPs)^14-19^. Cell-based methods could provide several potential advantages over current systems, including improved organ trafficking, tissue penetrance, cell-type specificity, transcriptional control for customizable gene programs, and decreased immunogenicity.

Previously published direct cell-to-cell gene editor delivery systems have employed immortalized cell lines, which would be difficult to translate clinically. Thus, adaptation to a primary human cell system is required for clinical feasibility. Given their ability to infiltrate tissues and their proliferative capacity – in addition to well-established FDA-approved pipelines for *ex vivo* modification – primary human T cells are ideal candidates for cell-based delivery of gene editors. Of the potential systems by which T cells could deliver gene editors, we reasoned that they may be well-suited to produce EDVs because T cells are the natural reservoir of human immunodeficiency virus (HIV), which serves as the backbone for several EDV systems. We therefore evaluated whether T cells could produce EDV architectures to package Cas9.

Here, we demonstrate that primary human T cells can produce a variety of retroviral-based EDVs that package functional Cas9 ribonucleoproteins (Cas9 EDVs). We term these engineered T cells Enveloped Particle Information Courier (EPIC-) T cells. We further demonstrate that primary human T cells can produce functional Cas9 EDVs pseudotyped with the fusogenic envelope protein Vesicular Stomatitis Virus Glycoprotein (VSV-G), which can transduce and edit an array of target cell types. We optimized lentiviral cassettes for EPIC-T cell generation and further developed a miniaturized Moloney leukemia virus (MLV) group-specific antigen polyprotein (Gag) that proved superior for T cell Cas9 EDV production. Finally, we designed and validated an all-in-one nanoplasmid for non-viral production of site-specific knock-in EPIC-T cells.

## RESULTS

### Primary human T cells can be engineered to produce diverse EDVs

To test the feasibility of EDV production by primary human T cells, we used plasmid nucleofection to test HIV and MLV-Gag derived EDV systems^16-18,20^ (Figure 1A). Beginning with the HIV-based system, we first collected supernatants to measure production of the HIV capsid protein P24 (Figure 1B) by ELISA. We detected robust levels of P24 in supernatants of T cells nucleofected with plasmids encoding HIV Gag-Pol alone or in combination with an HIV Gag-Cas9 fusion gene (Figure 1C). P24 levels were not affected by T cell stimulation with anti-CD3/CD28 dynabeads two days post-nucleofection. Levels of P24 secreted into the supernatant were consistent across T cells derived from multiple individuals (Figure 1D). Purified CD4 T cells and CD8 T cells produced similar levels as total unselected T cells (Figure 1E). P24 supernatant levels peaked three days after nucleofection before decaying, as expected with successive rounds of T cell proliferation and plasmid dilution (Figure S1A). Following nucleofection, there was no excess toxicity associated with T cells harboring packaging plasmids, as measured by flow cytometry (Figure S1B). We subsequently refer to primary human T cells engineered to produce EDVs as Enveloped Particle Information Courier (EPIC-) T cells.

**Figure 1.**
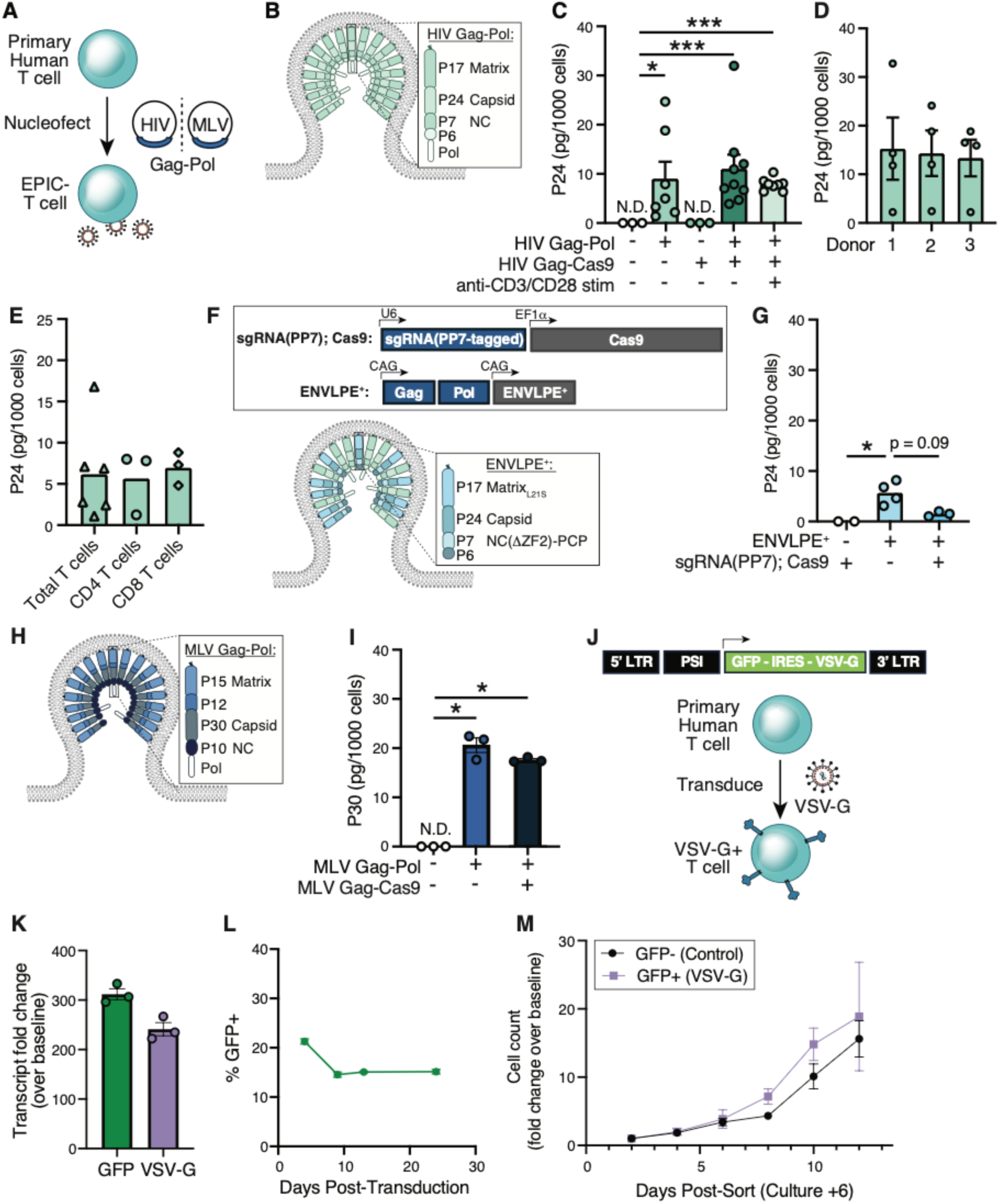
Primary human immune cells can produce a variety of enveloped delivery vehicles (EDVs). **(A)** Schematic for nucleofection of plasmids encoding HIV or MLV-derived Gag-Pol into primary human T cells to produce Enveloped Particle Information Courier-(EPIC-) T cells. **(B)** Schematic of HIV Gag on an EDV. **(C)** Levels of HIV P24 detected in the supernatant of primary human T cells two days after nucleofection with plasmids encoding the indicated genes, with and without post-nucleofection stimulation by anti-CD3/-CD28 dynabeads, normalized to T cell numbers. P-values from two-sided Student’s t-test. **(D)** Levels of HIV P24 detected in the supernatant of primary human T cells from three different donors nucleofected with HIV Gag-Pol plasmid, normalized to T cell numbers at time of nucleofection. **(E)** Levels of HIV P24 detected in the supernatant of primary human total T cells, CD4 T cells, or CD8 T cells nucleofected with plasmids encoding HIV Gag-Pol plasmid, normalized to cell numbers. **(F)** Schematics of (top) open reading frames for the ENVLPE^+^ system and (bottom) ENVLPE^+^ on an EDV. **(G)** Levels of HIV P24 detected in the supernatant of primary human T cells nucleofected with plasmid encoding ENVLPE^+^ alone or together with plasmid encoding sgRNA(PP7) and Cas9. P-values from two-sided Student’s t-test. **(H)** Schematic of MLV Gag on an EDV. **(I)** Levels of MLV P30 detected in the supernatant of primary human T cells nucleofected with plasmids encoding the indicated genes, normalized to T cell numbers. P-values from two-sided Student’s t-test. **(J)** Schematic of VSV-G construct with a GFP reporter. **(K)** Expression of GFP or VSV-G measured by qPCR from primary human T cells 10 days after transduction with lentivirus encoding the construct shown in (J). **(L)** Percent of primary human T cells expressing GFP over time after transduction with lentivirus encoding the construct shown in (J). **(M)** Number of GFP^+^ or GFP^-^ control sorted primary human T cells in culture over time after transduction with lentivirus encoding the construct shown in (J). Abbreviations: HIV = Human immunodeficiency virus. MLV = Moloney murine leukemia virus. Gag-Pol = Group-specific antigen and polymerase polyprotein. EPIC-T cell = Enveloped Particle Information Courier-T cell. EDV = enveloped delivery vehicle. ENVLPE = engineered nucleocytosolic vehicles for loading of programmable editors. SpCas9 = *Streptococcus pyogenes* Cas9. VSV-G = Vesicular Stomatitis Virus Glycoprotein. GFP = Green fluorescent protein. qPCR = quantitative polymerase chain reaction. * denotes nominal P-value < 0.05. ** denotes nominal P-value < 0.01.

Compared to HEK293T cells, the standard system used to produce EDVs *in vitro*, EPIC-T cells generated approximately 85-fold lower titers of P24 (Figure S1C). This difference may result from higher transfection rates of HEK293T cells as compared to nucleofection rates of primary T cells. We cloned an mCherry reporter into an HIV miniGag-Cas9 plasmid (∼10kb), approximately the same size as the HIV Gag-Pol packaging plasmid (10.7kb), to assess nucleofection rates of large plasmids. Four days following nucleofection, we detected mCherry expression in 11.9% of primary human T cells (Figure S1D). For comparison, after nucleofection of a 3.5kb GFP plasmid, we detected GFP expression in 28.1% of T cells. In contrast, after transfection with the larger HIV miniGag-Cas9 mCherry reporter plasmid, 93.6% of HEK293T cells exhibited mCherry fluorescence.

Two approaches by which cargo can be loaded into EDVs using retroviral Gag include through flexible protein linkers or an engineered Gag that can interact with aptamer-modified sgRNAs^19^. The latter creates an uncoupled architecture that could provide additional engineering flexibility in primary cells. We therefore tested an aptamer-based system, called engineered nucleocytosolic vehicles for loading of programmable editors (ENVLPE^+^)^19^. In this system, HIV Gag is modified with PP7 coat protein from Pseudomonas phage PP7, which binds to a PP7 aptamer-modified sgRNA complexed with Cas9 (Figure 1F). We detected P24 in the supernatant of T cells nucleofected with ENVLPE^+^ constructs (Figure 1G). However, in contrast to HIV Gag-Pol (Figure 1C), addition of a second plasmid encoding a PP7-tagged sgRNA cassette and Cas9 during nucleofection with ENVLPE^+^ was associated with an approximately two-fold reduction in detectable P24 titer (Figure 1G), which may result from nucleofection toxicity due to larger plasmid sizes (14.2kb for ENVLPE^+^ vs. 10.7kb for HIV Gag-Pol).

We next assessed whether EDVs based on a different retrovirus architecture could also be produced by primary human T cells. We nucleofected primary human T cells with plasmids expressing Gag-Pol from Moloney murine leukemia virus (MLV) and observed detectable P30 capsid protein at levels that were consistently higher on a per-cell basis than those we had observed for HIV P24 (Figure 1H-I). Concordantly, the difference in levels of P30 in nucleofected T cell supernatants versus HEK293T cell supernatants was also smaller (18.1-fold) (Figure S1E).

To facilitate cargo delivery by EPIC-T cell-produced EDVs, we next engineered a fusogenic envelope protein into the EDVs. The most commonly used fusogenic envelope protein is vesicular stomatitis virus (VSV) glycoprotein (VSV-G), and VSV-infected T cells have been shown to produce infectious virions^21^, suggesting they should be capable of expressing VSV-G. We therefore cloned a bicistronic expression cassette linking GFP expression to VSV-G expression and used it to transduce primary human T cells (Figure 1J). Following transduction, we detected substantial expression of both GFP and VSV-G RNA transcripts by qPCR (Figure 1K) and VSV-G protein expression by Western blot (Figure S1F). GFP-positive cells displayed only marginally higher rates of cell death compared to GFP-negative cells (3% vs. 1%) (Figure S1G). Over time, the percent of GFP-positive T cells remained stable (Figure 1L), and sorted GFP-positive cells displayed equivalent rates of proliferation compared to GFP-negative controls (Figure 1M). However, we detected decreases in VSV-G transcript, GFP transcript, and GFP median fluorescence intensity over time in culture, suggesting some degree of epigenetic silencing may occur (Figure S1H-I).

### Primary human T cells can produce functional Cas9 EDVs that edit target cells

Having established that EPIC-T cells could produce retroviral-based EDVs, we next sought to confirm whether T cell EDVs could package Cas9 nuclease from *Streptococcus pyogenes* (Cas9). Among gene editors, we chose Cas9 as an initial cargo in part due to its large size (1368 amino acids) that poses stoichiometric constraints^20^, reasoning that, if EPIC-T cells could produce functional Cas9 EDVs, then they would also likely be able to package other cargo. Analogous to HEK293T cells, which cannot produce full-length HIV Gag-Cas9 EDVs^16^, nucleofection of HIV Gag-Cas9 into primary human T cells resulted in minimal detectable Cas9 protein in the supernatant as detected by ELISA, but levels increased significantly when T cells were co-electroporated with Gag-Pol (which together constitute the VLPv2 system^16^) (Figure 2A-B). In the latter condition, Cas9 protein in the supernatant was also detected by Western blot (Figure S2A). In contrast to full-length Gag-Cas9, nucleofection of T cells with a plasmid encoding a minimized (mini)Gag-Cas9 fusion gene^15^ resulted in higher Cas9 protein levels in the supernatant even without Gag-Pol (Figures 2B, S2B-C), suggesting that EPIC-T cells can generate minimized EDVs (miniEDVs). Compared to transfected HEK293T cells, EPIC-T cells produced 14.3- and 10.4-fold less Cas9 protein in the supernatant for VLPv2 and miniEDV architectures, respectively (Figure S2C).

**Figure 2.**
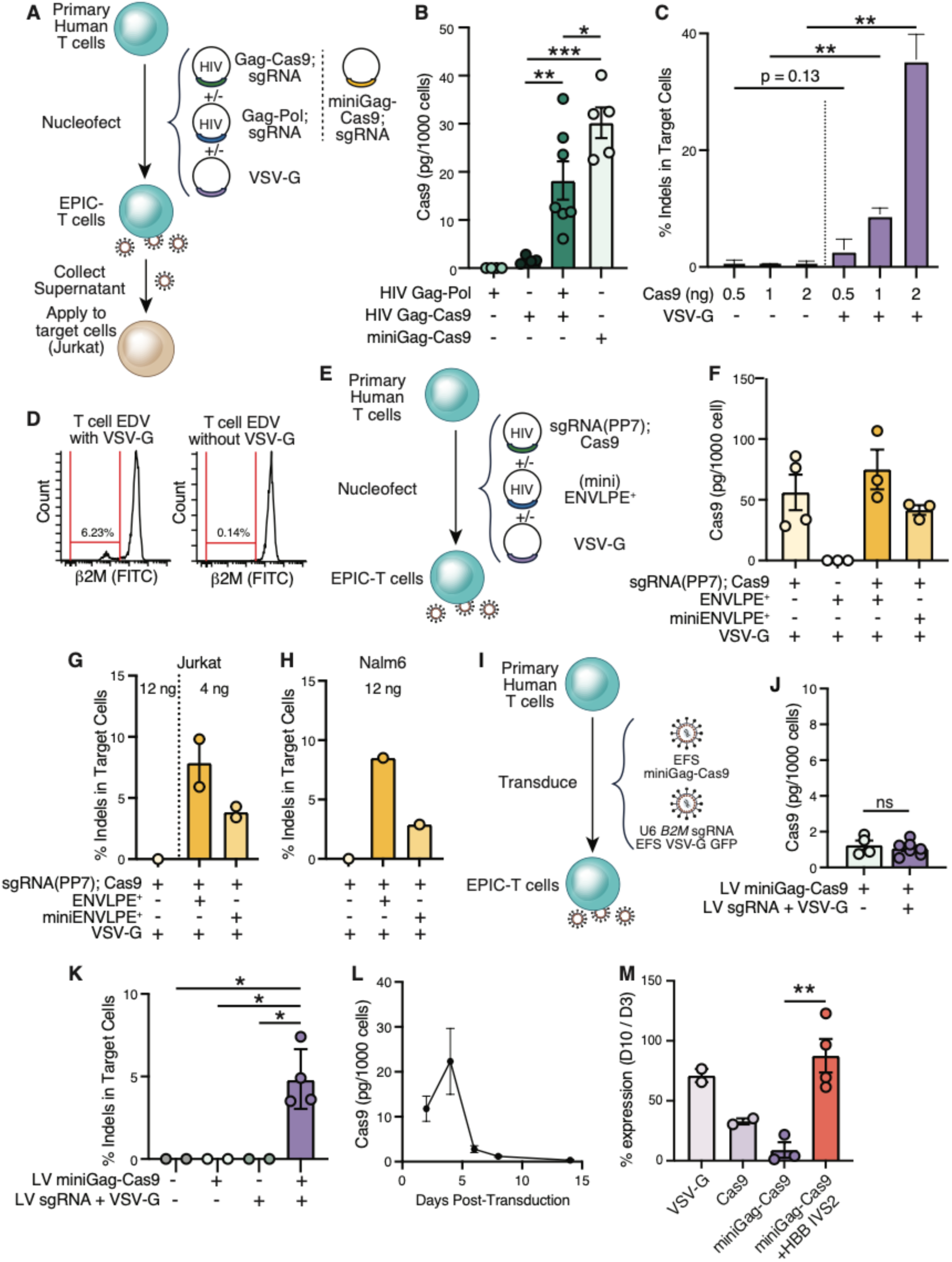
T cells can produce EDVs that package functional Cas9 ribonucleoproteins capable of gene editing in target cells. **(A)** Schematic for nucleofection of plasmids encoding Gag-Pol, Gag-Cas9, and/or VSV-G into primary human T cells to produce EPIC-T cells. Supernatants from EPIC-T cells were collected and transferred onto target cells (Jurkat) to assess gene editing. **(B)** Levels of Cas9 detected in the supernatant of primary human T cells nucleofected with plasmids encoding the indicated genes, normalized to T cell numbers. P-values from two-sided Student’s t-test. **(C)** Percent of *B2M* gene amplicons with indels in Jurkat target cells incubated with supernatants from primary human T cells nucleofected with plasmids encoding HIV Gag-Pol and Gag-Cas9, with or without plasmid encoding VSV-G. Cas9 levels in the supernatant were quantified by ELISA and used to calculate the quantity of supernatant transferred as indicated. P-values from two-sided Student’s t-test. **(D)** Flow cytometry analysis of β2M surface protein expression on target Jurkat cells incubated with supernatants from primary human T cells nucleofected with plasmids encoding HIV Gag-Pol and Gag-Cas9, with (left) or without (right) plasmid encoding VSV-G. **(E)** Schematic for nucleofection of plasmids encoding (mini)ENVLPE^+^, sgRNA(PP7), and/or VSV-G into primary human T cells to produce EPIC-T cells. **(F)** Levels of Cas9 detected in the supernatant of primary human T cells nucleofected with plasmids encoding the indicated genes, normalized to T cell numbers. **(G-H)** Percent of *B2M* gene amplicons with indels in (G) Jurkat target cells or in (H) Nalm-6 target cells incubated with supernatants from primary human T cells nucleofected with plasmids encoding the indicated genes. **(I)** Schematic of EPIC-T cells produced using a lentivirus-based approach. Primary human T cells were transduced with lentivirus encoding miniGag-Cas9 and a second lentivirus encoding sgRNA targeting *B2M* along with VSV-G and GFP. **(J)** Levels of Cas9 detected in the supernatant of primary human T cells transduced with lentiviruses encoding the indicated genes, normalized to T cell numbers. P-value from two-sided Student’s t-test. **(K)** Percent of *B2M* gene amplicons with indels in Jurkat target cells incubated with supernatants from primary human T cells transduced with lentiviruses encoding the indicated genes. P-values from two-sided Student’s t-test. **(L)** Levels of Cas9 detected over time in the supernatant of primary human T cells transduced with lentiviruses encoding miniGag-Cas9 and sgRNA + VSV-G, normalized to T cell numbers. **(M)** Flow cytometry analysis of bicistronic marker percent expression (GFP for VSV-G and Cas9, surface expressed RQR8 for miniGag-Cas9 with and without HBB IVS) in primary human T cells on day 10 after transduction, normalized to percentage on day 3 after transduction. T cells were transduced with lentivirus encoding the indicated genes. Abbreviations: Gag-Pol = Group-specific antigen and polymerase polyprotein. Gag-Cas9 = Group-specific antigen Cas9 fusion gene. VSV-G = Vesicular Stomatitis Virus Glycoprotein. EPIC-T cell = Enveloped Particle Information Courier-T cell. *B2M* = Beta-2-microglobulin gene. indel = insertion or deletion mutation. β2M = Beta-2-microglobulin protein. miniGag = minimized HIV Group-specific antigen. sgRNA = single guide ribonucleic acid. GFP = Green fluorescent protein. ENVLPE = engineered nucleocytosolic vehicles for loading of programmable editors. * denotes nominal P-value < 0.05. ** denotes nominal P-value < 0.01.

ELISA detection of Cas9 in supernatants does not ensure its functionality. Moreover, recent studies have shown that T cells can produce exosomes that can be taken up by tumor cells^22^. Therefore, it was unclear whether T cell-derived EDVs produced in the absence of any pseudotyping envelope protein could deliver cargo to target cells. To determine whether T cells produced functional Cas9 EDVs, we nucleofected T cells with VLPv2 plasmids containing an sgRNA targeting the beta-2-microglobulin gene (*B2M*) with and without VSV-G (Figure 2A). We used ELISA to quantify Cas9 concentration in EPIC-T cell supernatant and applied volume-normalized doses to target Jurkat cells (Figure 2A). Three days following the addition of supernatant, we assessed indel formation at the target *B2M* locus in treated Jurkat cells by amplicon sequencing and found that indel formation was dose responsive and VSV-G dependent (Figure 2C). Moreover, a VSV-G-dependent population of Jurkat cells lacking β2M surface protein expression was evident by flow cytometry (Figure 2D). When normalized to levels of Cas9 protein, EPIC-T cell supernatants and HEK293T cell supernatants performed similarly in their ability to create indels in target Jurkat cells (Figure 2C, S2D).

We further tested whether primary human T cells could produce functional Cas9-containing EDVs using the ENVLPE^+^ system (Figure 2E). We detected Cas9 protein in the supernatant of T cells nucleofected with a plasmid encoding both PP7-modified *B2M* sgRNA and Cas9 even without retroviral Gag (Figure 2F). Indeed, while co-nucleofection of plasmids encoding either ENVLPE^+^ or miniENVLPE achieved detectable levels of Cas9 in T cell supernatants, levels were similar to those achieved by nucleofection of only the plasmid encoding Cas9 (Figure 2F). However, we only detected *B2M* indels in target Jurkat and Nalm6 cells when adding supernatants from T cells co-nucleofected with plasmids encoding ENVLPE^+^ or miniENVLPE EDV packaging systems, but not when adding supernatant from T cells nucleofected with plasmids encoding Cas9, VSV-G, and sgRNA (Figure 2G-H), even at higher doses than those previously tested (Figure S2E). This result indicated that Cas9 was not functionally delivered to target cells without EDV machinery, even when it was detectable in T cell supernatants.

### Development of a lentivirus-based approach to produce EPIC-T cells

A major advantage of cell therapies is their ability to proliferate *in vivo*, amplifying a desired effect. Unfortunately, this benefit is negated by plasmid nucleofection due to plasmid dilution during T cell proliferation; additionally, plasmid nucleofection can result in substantial toxicity^23^. To address these challenges, we developed a dual lentivirus system to integrate EDV machinery into primary human T cells (Figure 2I). One lentivirus encodes a U6 promoter-driven sgRNA cassette and VSV-G envelope protein, while the other encodes miniGag-Cas9 to drive miniEDV production. EDV systems that require expression of Gag-Pol, such as VLPv2, are disfavored for therapeutic applications due to safety considerations and their prohibitively large size. Minimized Gag does not contain nucleocapsid (Figure S2B), which natively binds viral RNA, and therefore should not package viral transcript^24^. Indeed, miniEDVs with and without Cas9 cargo, when co-transfected with a control lentiviral transfer plasmid containing GFP, packaged minimally detectable lentiviral transcript (Figure S2F-G). When HIV Rev was deleted from the original miniGag system, levels of detectable transcript further decreased (Figure S2G). Moreover, given the absence of Pol, which encodes for reverse transcriptase and integrase, there was no successful propagation of transfer plasmid by miniEDVs (Figure S2H).

EDV expression plasmids utilize a strong synthetic CAG promoter, which does not translate well to lentiviral systems due to large size and high GC content interfering with viral RNA packaging. Thus, we screened for alternative promoters for miniGag-Cas9. Both HEK293T and EPIC-T cell miniEDV production, measured by Cas9 protein levels in supernatants, was highly promoter dependent (Figure S2I). There was no significant difference among the alternative promoters tested, and we selected the most compact promoter, Elongation Factor 1-alpha short (EFS), for further testing (Figure 2I). After transduction with lentivirus encoding miniGag-Cas9 alone or together with lentivirus encoding sgRNA and VSV-G, EPIC-T cells produced detectable Cas9 protein in supernatant at levels similar to those observed for EPIC-T cells made by plasmid nucleofection when using the same promoter (Figure 2J, S2I). Supernatant collected from single-transduced T cells expressing either miniGag-Cas9 or from single-transduced T cells expressing U6-driven *B2M* sgRNA and VSV-G did not generate indels in target Jurkat cells, as expected (Figure 2K). However, supernatant from double-transduced, EPIC-T cells successfully edited target cells (Figure 2K). Optimal lentiviral preparations achieve approximately 70-80% transduction efficiency in primary human T cells depending on the size of the desired integration and the health of the T cell population^25^. In this study, the lentivirus encoding sgRNA and VSV-G routinely achieved 50-70% transduction efficiency, while the lentivirus encoding miniGag-Cas9 typically achieved only 5-8% efficiency. Co-transduction of T cells thus produced a mixed culture of untransduced T cells, single-transduced T cells expressing either miniGag-Cas9 or sgRNA and VSV-G, and double-transduced EPIC-T cells expressing all components for EDV generation: miniGag-Cas9, sgRNA, and VSV-G. In this culture system, we observed increasing levels of *B2M* knockout cells over time, likely from EPIC-T cells editing singly transduced and untransduced T cell neighbors (Figure S2J). In contrast, expression of GFP (encoded by the lentivirus containing sgRNA and VSV-G) proportionally decreased over time (Figure S2K). This observation provides further support that EPIC-T cells can package functional miniEDVs that edit non-producer, target cells.

While characterizing these EPIC-T cell cultures, we noticed that levels of Cas9 protein in the supernatant continued to decrease over time, peaking at day 4 post transduction before rapidly declining to nearly undetectable levels by day 8 (Figure 2L). Because we did not observe substantial cell loss among sorted T cells that were singly transduced with miniGag-Cas9 (Figure S2L), and due to the known role of HUSH1 in silencing intronless segments >1kb, such as those present in lentivirus^26-28^, we hypothesized that this result was due to epigenetic silencing rather than toxicity. We cloned a bicistronic cell surface marker (RQR8^29^) into the miniGag-Cas9 transfer cassette and detected complete loss of marker expression by day 14 post transduction (Figure S2M-N). We modified the construct so that separate promoters controlled miniGag-Cas9 (5307 bp) and RQR8 (474 bp) expression and detected no loss of RQR8 expression at day 14 (Figure S2M-N). These results are consistent with a role for epigenetic silencing in the decline of detectable Cas9 protein in EPIC-T cell supernatants over time. Of note, we also observed decreasing levels of a bicistronic GFP marker expressed in combination with either VSV-G or Cas9 (Figure 2M). Likely due to their smaller size, the loss of marker expression was not as pronounced by day 10 as with miniGag-Cas9. A recent study found that Cas9 transgene silencing could be mitigated by including introns, such as the 850bp second intervening sequence (intron 2; IVS2) of the human beta-globin (HBB) gene (HBB IVS2)^26,27^. Consistent with this finding, inclusion of HBB IVS2 in the 5’ UTR largely mitigated bicistronic marker loss at day 10 and was subsequently included in our lentiviral construct designs (Figure 2M).

The lentiviruses used to transduce primary human T cells are produced by HEK293T producer cells. Because the miniGag-Cas9 fusion gene is encoded on the transfer plasmid transfected into HEK293T cells, and because miniGag-Cas9 retains C-terminal capsid Gag-Gag interaction domains, we reasoned that HEK293T lentivirus producer cells would generate mosaic lentivirus composed of some percentage of wild-type Gag and some percentage of miniGag-Cas9 (Figure S2O). It was unclear whether such a mosaic lentivirus would exhibit impaired infectivity, which could limit the production of EPIC-T cells. To address this, we generated lentiviral particles of varying mosaicism by co-transfecting different ratios of psPAX2 packaging plasmid (encoding wild-type HIV Gag-Pol) with plasmids encoding miniGag or miniGag-Cas9, VSV-G, and a control lentiviral transfer plasmid encoding for CD19 and GFP. Increasing miniGag mosaicism decreased viral infectivity but still achieved near maximal transduction at the highest doses tested (Figure S2P). In contrast, increasing miniGag-Cas9 mosaicism drastically reduced viral infectivity, which was only partially ameliorated by increasing the dose (Figure S2P). Decreased infectivity could be explained in part by decreased viral transcript packaging by mosaic particles (Figure S2Q). Based on these data, we expected the lentivirus encoding miniGag-Cas9 to exhibit decreased infectivity. Indeed, compared to lentivirus encoding either miniGag or a control transfer plasmid, miniGag-Cas9 lentivirus exhibited impaired transduction (Figure S2R).

To approximate the degree of mosaicism, we measured P24 and Cas9 levels by ELISA from lentiviral preparations of transfer plasmids containing miniGag-Cas9. We interpreted an increased P24/Cas9 ratio as being less mosaic and therefore likely more effective lentivirus. Our studies had utilized third-generation transfer plasmids, which have an upstream CMV promoter that drives high levels of viral transcript. We wondered whether expression in a second-generation transfer plasmid, with a comparatively weaker upstream 5’ LTR promoter driving viral transcript, could improve P24/Cas9 ratios. Downstream in a transfer plasmid, a second promoter drives expression of the transgene insert. P24/Cas9 ratios were equivalent between second- and third-generation transfer plasmids encoding miniGag-Cas9 when the transgene insert was driven by a downstream constitutive EFS promoter (Figure S2S). However, when the transgene insert was driven by a downstream inducible UAS CMV promoter, second-generation transfer plasmid indeed displayed an improved P24/Cas9 ratio.

### Development of a miniature MLV-Gag EDV

Despite these improvements, decreased infectivity of the lentivirus encoding miniGag-Cas9 remained a severe bottleneck for EPIC T-cell characterization and functional studies. Given our earlier observation that T cells produced higher titers of MLV P30 than HIV P24 on a per-cell basis (Figure 1C, 1I), we next aimed to identify a MLV Gag-Cas9 variant that produced Cas9 EDVs, ideally without requiring MLV Gag-Pol, and that was compact enough to fit in a lentiviral transfer system.

In HEK293T cells, we observed that transfection of plasmid encoding MLV Gag-Cas9 resulted in high levels of detectable Cas9 protein in the supernatant, which were not further increased by addition of MLV Gag-Pol (Figure 3A-B). These results suggest that MLV Gag-Cas9, in contrast to HIV Gag-Cas9, is sufficient for Cas9 EDV generation. To confirm that Cas9 was functionally packaged, we transfected HEK293T cells with plasmids encoding MLV Gag-Cas9, VSV-G, and *B2M* sgRNA without MLV Gag-Pol and applied the collected supernatant to target HEK293T or Jurkat cells. In a VSV-G dependent manner, MLV Gag-Cas9 EDVs elicited indels in both types of target cells (Figure 3C). Similar to our prior controls, supernatant from cells transfected with plasmids encoding Cas9, *B2M* sgRNA, and VSV-G did not produce indels in target cells (Figure 3C).

**Figure 3.**
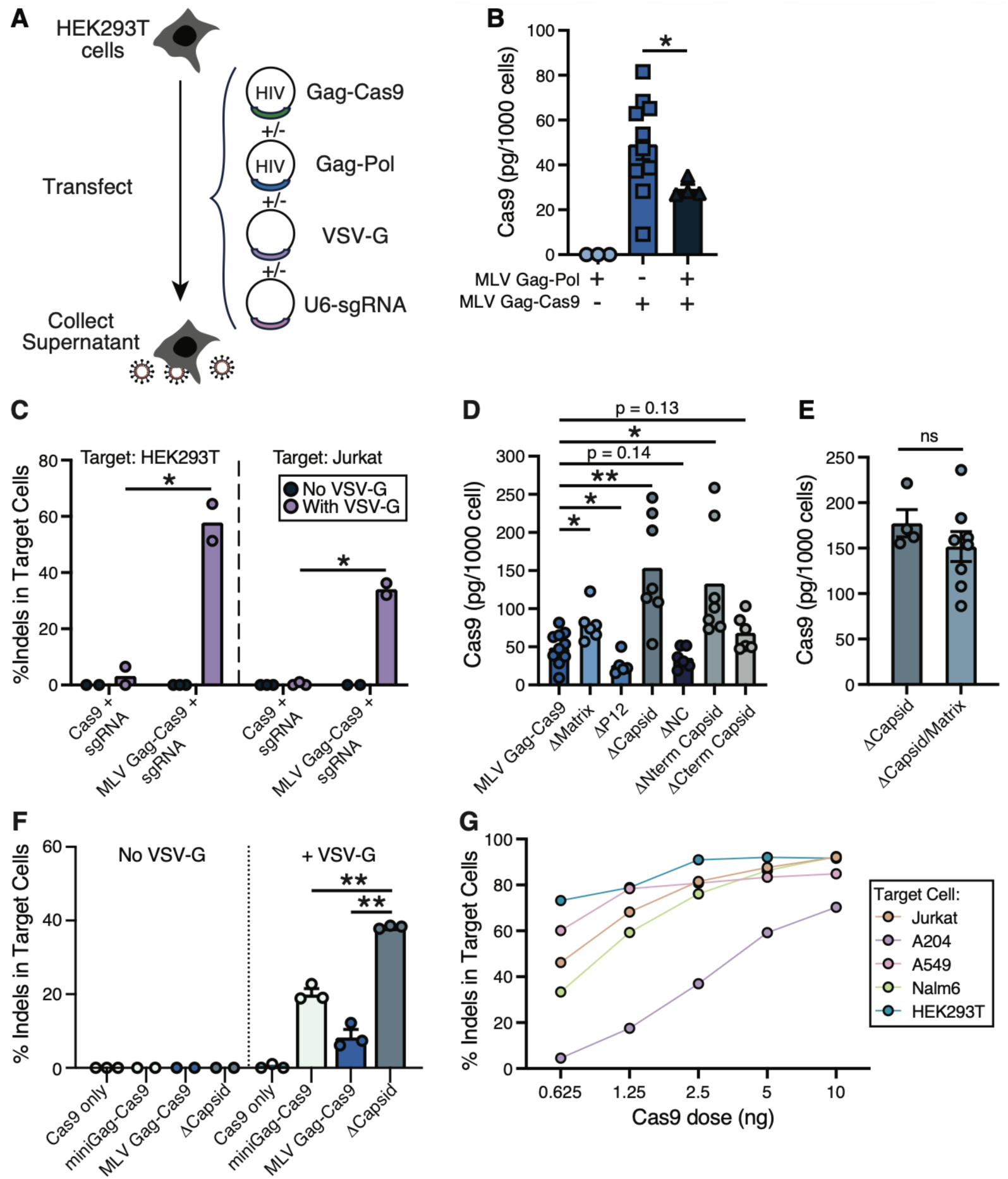
Development of a miniaturized MLV-based EDV system. **(A)** Schematic for transfection of plasmids encoding MLV Gag-Pol, Gag-Cas9, U6-sgRNA, and/or VSV-G into HEK293T cells. **(B)** Levels of Cas9 detected in the supernatant of HEK293T cells transfected with plasmids encoding the indicated genes, normalized to cell numbers. P-values from two-sided Student’s t-test. **(C)** Percent of *B2M* gene amplicons with indels in HEK293T (left) or Jurkat (right) target cells incubated with supernatants from HEK293T cells transfected with plasmids encoding Cas9 only or MLV Gag-Cas9 fusion, with or without plasmid encoding VSV-G. **(D-E)** Levels of Cas9 detected in the supernatant of HEK293T cells transfected with plasmids encoding the indicated genes, normalized to cell numbers. Δ indicates deletion of the indicated component of MLV Gag from the MLV Gag-Cas9 fusion gene. P-values from two-sided Student’s t-test. **(F)** Percent of *B2M* gene amplicons with indels in Jurkat target cells incubated with supernatants from HEK293T cells transfected with plasmids encoding the indicated genes, with or without plasmid encoding VSV-G. **(G)** Percent of *B2M* gene amplicons with indels in different target cell types incubated with varying supernatant doses from HEK293T cells transfected with plasmids encoding U6-*B2M* sgRNA, MLV GagΔCapsid-Cas9 and VSV-G. Abbreviations: Gag-Pol = Group-specific antigen and polymerase polyprotein. Gag-Cas9 = Group-specific antigen Cas9 fusion gene. VSV-G = Vesicular Stomatitis Virus Glycoprotein. EPIC-T cell = Enveloped Particle Information Courier-T cell. *B2M* = Beta-2-microglobulin gene. indel = insertion or deletion mutation. miniGag = miniaturized HIV Group-specific antigen. * denotes nominal P-value < 0.05. ** denotes nominal P-value < 0.01. *** denotes nominal P-value < 0.005.

We next explored whether MLV Gag-Cas9 could be miniaturized to improve lentiviral compatibility. We generated a series of MLV Gag-Cas9 mutants with individual subunit deletions and screened transfected HEK293T supernatants for Cas9 levels. Among the tested mutants, deletion (Δ) of matrix, capsid, or N-terminal capsid resulted in increased levels of Cas9 protein in supernatants, while deletion of p12 had the opposite effect, likely due to its known role in recruiting ESCRT machinery for viral budding^30^ (Figure 3D). Deletion of the MLV Gag myristoylation domain (MLV Gag(ΔMyris)), which allows anchoring to the cell membrane, resulted in nearly the same Cas9 supernatant levels as unlinked Cas9, suggesting a pathway of egress for Cas9 apart from EDV packaging (Figure S3A). Neither co-transfection with MLV Gag-Pol nor additive truncations, MLV Gag(ΔMatrix) plus MLV Gag(ΔCapsid), further increased levels of Cas9 protein in supernatant (Figure 3E, S3B).

When normalizing for Cas9 levels, MLV Gag(ΔCapsid)-Cas9 EDVs maintained functionality that was equivalent to HIV miniGag-Cas9 miniEDVs and VLPv2 EDVs at saturating doses against Jurkat targets (Figure S3C). However, MLV Gag(ΔCapsid)-Cas9 EDVs proved to be superior at editing Jurkat target cells at a lower dose (Figure 3F). In contrast to these results, despite similarly increasing levels of Cas9 protein in the supernatant, MLV Gag(ΔMatrix)-Cas9 EDVs exhibited drastically reduced target cell editing, indicating that matrix was critical for generation of functional EDVs (Figure S3D). Similarly, when supernatant from Gag(ΔMyris)-Cas9 EDVs was applied to target cells, minimal indels were detected even in the presence of VSV-G (Figure S3D). Given that different cell types exhibit distinct susceptibilities to different retroviruses and envelope proteins, we tested the ability of MLV Gag(ΔCapsid)-Cas9 EDVs to generate indels in a variety of additional cell types including A204, Nalm6, and A549 cells, and we successfully detected indels in all tested target cell types (Figure 3G).

Arising from our initial interest in adapting lentivirus for integration of EDV production machinery, we co-transfected HEK293T cells with GFP transfer plasmid, envelope, and equimolar amounts of HIV Gag-Pol and either MLV Gag-Cas9 or MLV GagΔCapsid-Cas9. Both MLV Gag-Cas9 and MLV GagΔCapsid-Cas9 decreased infectivity compared to HIV Gag-Pol control (Figure S3E). However, the decrease was mild compared to miniGag-Cas9 (Figure S2P), suggesting either decreased mosaicism due to fewer interactions between HIV and MLV Gag or improved infectivity of a mosaic particle. In summary, we identified that MLV Gag-Cas9 was sufficient for functional EDV particle formation based on ELISA and editing of target cells. Furthermore, we engineered a superior particle by deleting MLV Capsid, which resulted in higher levels of Cas9 protein in supernatants and more efficient editing of target cells.

### T cells can produce miniaturized MLV EDVs

Our primary interest remained translation of this miniaturized MLV EDV system into primary human T cells. To simplify EDV generation, we cloned a *B2M* sgRNA driven by a U6 promoter into the MLV Gag(ΔCapsid)-Cas9 expression plasmid, which was co-nucleofected with a plasmid encoding VSV-G to generate EPIC-T cells (Figure 4A). Following plasmid nucleofection, we detected higher levels of Cas9 protein in supernatants of EPIC-T cells producing MLV Gag(ΔCapsid)-Cas9 EDVs compared to those producing MLV Gag-Cas9 EDVs (Figure 4B). Expression of MLV Gag(ΔCapsid)-Cas9, like other EDV-generating constructs, is expected to generate indels within the producer cell, and detection of self-knockout among producer cell populations can confirm proper expression^16,20^. As expected, we detected indels at the *B2M* locus in T cells following introduction of EDV plasmids, and levels were similar to those in T cells nucleofected with only a Cas9 expression plasmid and sgRNA (Figure S4A). To explore the versatility of T cell production of MLV Gag(ΔCapsid)-Cas9 EDVs, we linked it to *Staphylococcus aureus* Cas9 (saCas9) and similarly observed significant levels of saCas9 protein in the supernatant (Figure S4B). Once more, MLV Gag(ΔCapsid)-saCas9 resulted in higher detected levels of saCas9 in supernatants compared to MLV Gag-saCas9.

**Figure 4.**
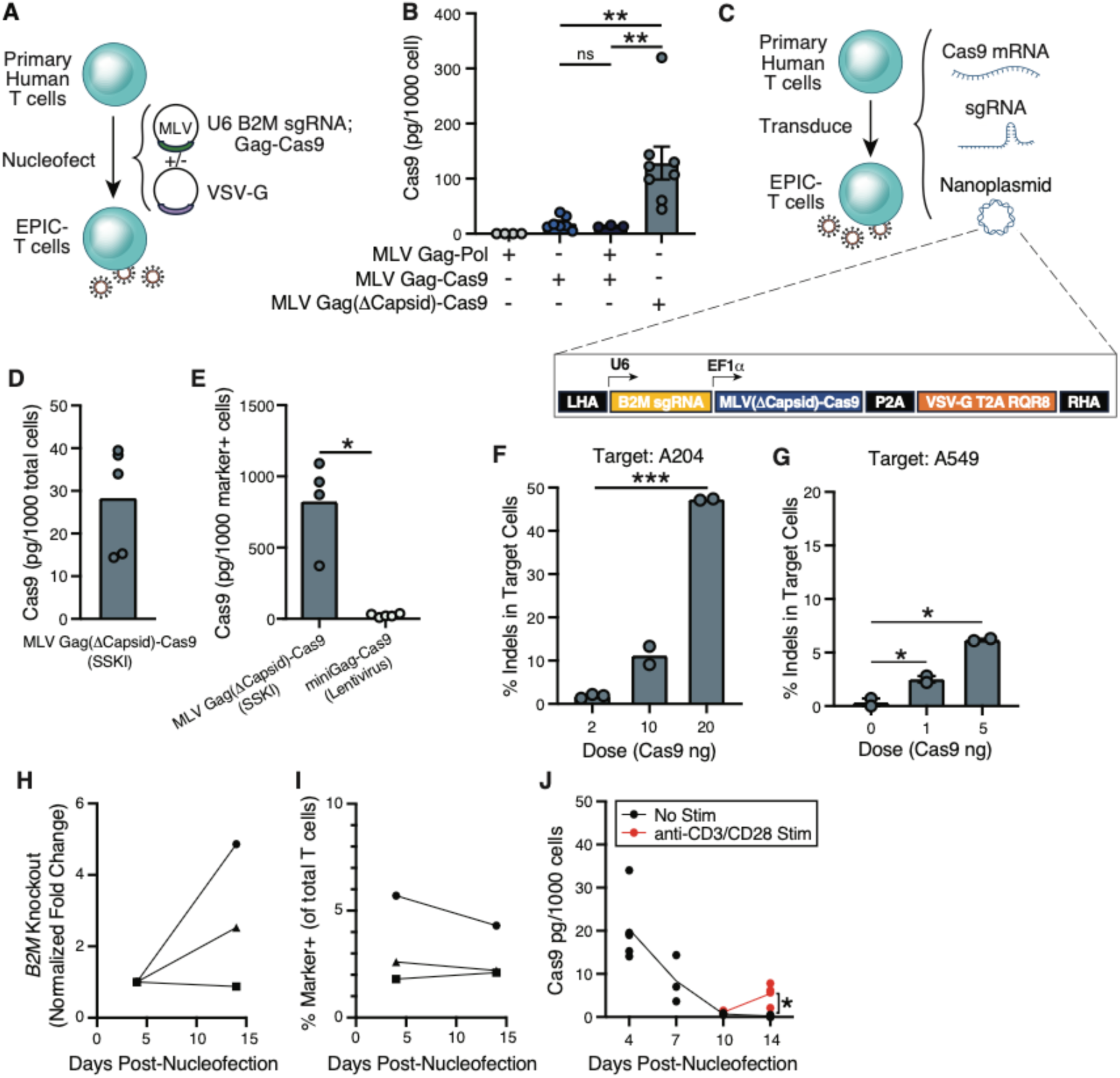
Generation of EPIC T cells via all-in-one, site-specific knock-in (SSKI) of miniaturized MLV EDV production machinery. **(A**) Schematic for nucleofection of plasmids encoding MLV Gag-Cas9 into primary human T cells to produce EPIC-T cells. **(B)** Levels of Cas9 detected in the supernatant of primary human T cells nucleofected with plasmids encoding the indicated genes, normalized to cell numbers. Δ indicates deletion of the indicated component of MLV Gag. P-values from two-sided Student’s t-test. **(C)** Schematic for SSKI of an all-in-one double-stranded DNA template encoding U6-sgRNA, MLV Gag(ΔCapsid)-Cas9 fusion gene, and VSV-G into primary human T cells to produce EPIC-T cells. **(D)** Levels of Cas9 detected in the supernatant of primary human EPIC-T cells generated by SSKI, normalized to total cell numbers. **(E)** Levels of Cas9 detected in the supernatant of primary human EPIC-T cells generated by SSKI of MLV GagΔCapsid-Cas9 or by lentiviral transduction of miniGag-Cas9, normalized to marker^+^ cell numbers. **(F-G)** Percent of *B2M* gene amplicons with indels in **(F)** HER2^+^ A204 target cells and **(G)** A549 and incubated with supernatants from primary human EPIC-T cells generated by SSKI. **(H)** Fold-change in expression of surface *B2M* in SSKI EPIC-T cell cultures over time measured by flow cytometry. **(I)** Percent of marker^+^ T cells in SSKI EPIC-T cell cultures over time. **(J)** Levels of Cas9 detected in the supernatant of SSKI EPIC-T cells normalized to total cell numbers over time with (red line) or without (black line) anti-CD3/CD28 dynabeads (1:1) added at day 8 post nucleofection. Abbreviations: MLV Gag-Cas9 = Moloney murine leukemia virus Group-specific antigen-Cas9 fusion gene. EPIC-T cell = Enveloped Particle Information Courier-T cell. VSV-G = Vesicular Stomatitis Virus Glycoprotein. *B2M* = Beta-2-microglobulin gene. indel = insertion or deletion mutation. * denotes nominal P-value < 0.05. ** denotes nominal P-value < 0.01.

### Non-viral EPIC-T cell production by site-specific knock-in

While lentiviral manufacturing of T cells for cell therapy remains the current standard, non-viral engineering strategies have several advantages. Site-specific knock-in is less prone to epigenetic silencing across cell divisions, enables more precise control of transgene expression, does not risk oncogenic or multi-integration events, and has potential advantages in terms of production cost^31^. We therefore explored site-specific knock-in (SSKI) of a nanoplasmid-based homology-directed repair template (HDRT) as an alternative, non-viral strategy to engineer EPIC-T cells (Figure 4C). The size of EDV production machinery (U6 – sgRNA, MLV GagΔCapsid-Cas9, and VSV-G) approaches 10kb, the published limit of HDRT knock-ins^32,33^. Therefore, we took advantage of the recent optimization of ultra large site-specific knock-ins^33^, and we employed Cas9 mRNA and a compact nanoplasmid to knock-in EDV production machinery into an open region on Chromosome 11 in primary human T cells. Because HUSH1 silencing can still occur at open chromatin regions^27^, we included HBB IVS2 to mitigate silencing of the knock-in cassette.

Following knock-in, we observed production of Cas9 protein in the supernatant (Figure 4D). Because our nanoplasmid encodes a cell surface marker (RQR8), we normalized Cas9 levels to the percent of cells expressing the marker, resulting in estimated secretion of Cas9 protein levels exceeding 0.7 picograms/cell, which far exceeded per-cell levels detected from lentiviral introduction of miniGag-Cas9 (Figure 4E). Transfer of knock-in EPIC-T cell supernatant successfully generated indels in target A549 and A204 cells (Figure 4F-G). Moreover, we noted an increase in *B2M* knockout cells among our T cell cultures over time, consistent with EPIC-T cell transfer of Cas9 EDVs to other T cells in culture (Figure 4H). The percent of T cells expressing the RQR8 marker remained stable over time, demonstrating stability of the construct expression (Figures 4I, S4C). Importantly, despite stable marker expression, we still observed a decrease in Cas9 protein levels detected in supernatants over time when normalized for cell numbers (Figure 4J). Because primary T cells have low resting transcriptional activity that is rapidly upregulated upon activation^34,35^, we reasoned that stimulation may promote EDV production in this context. Therefore, eight days following nucleofection, when levels of Cas9 in the supernatant declined, we provided EPIC T-cells with anti-CD3/CD28 stimulation and observed a significant increase in Cas9 protein levels in the supernatant six days later (Figure 4J) despite similar total cell numbers (Figure S4D). These results indicate that EDV production by EPIC T-cells produced by non-viral knock-in is dynamically regulated as a function of their activation state, which has translational implications.

In this manuscript, we demonstrate that primary human T cells can generate retroviral Gag-based EDVs that package Cas9 ribonucleoproteins. Moreover, we show that T cell-generated EDVs can be pseudotyped with VSV-G and transduce a variety of target cell populations. Among published EDV architectures, T cells were particularly amenable to production of MLV-based EDVs. Given this, we engineered a new miniaturized MLV Gag EDV system that demonstrated superiority compared to existing frameworks and could be site-specifically integrated into T cells. We term these EDV-producing T cells as Enveloped Particle Information Courier-(EPIC-) T cells to reflect the membrane derivation of EDVs, the ability to load EDVs with varying protein and RNA cargo, and the cell-based approach. We anticipate that cellular delivery of gene editing machinery will have broad translational applications to a variety of diseases.

## DISCUSSION

An EDV is composed of a membrane-associated assembly protein capable of vesicle budding and a fusogenic envelope protein which allows entry into target cells. Here, we explored the ability of primary human T cells to generate EDVs derived from two assembly proteins (HIV and MLV Gag) packaging Cas9 ribonucleoproteins and pseudotyped by a single envelope protein (VSV-G). We also expect T cells to be able to produce additional EDV architectures, package alternative cargoes, and produce EDVs pseudotyped by additional envelope proteins beyond those tested here. It is possible that alternative retroviral Gag proteins or membrane assembly systems not tested may prove superior for T cell EDV production. It is also likely that the optimal EDV production system may be specific to the desired cargo. Different approaches may prove more adept depending on whether the cargo is RNA or protein, the specific cargo loading mechanisms, and the cargo stoichiometry.

Engineering of primary T cells to produce EDVs proved challenging due to epigenetic silencing and the ultra-large sequences needed for Cas9 EDV production. To overcome these obstacles, we validated a variety of approaches to generate EPIC-T cells, including an all-in-one site-specific knock-in system, which proved superior to lentivirus or nucleofection strategies. Depending on the context, it is also possible that alternative primary human immune or non-immune cell types may be amenable to cellular EDV production. Given the breadth of cells readily infected by and capable of producing retrovirus, we posit that additional cell types will be capable of enveloped particle information couriering, including myeloid cells.

In the future, we envision pairing EPIC-T cell technology with additional T cell engineering approaches to maximize their ability to edit target cells *in vivo* with high specificity. For instance, custom synthetic receptors like SNIPRs^36^ enable bespoke transcriptional control of payloads, which could include EDV production machinery. Co-expression of chimeric antigen receptors (CAR) or transgenic T cell receptors (TCRs) in EPIC-T cells could direct target cell specificity and promote EPIC-T cell proliferation. Moreover, overexpression of chemokine receptors or adhesion molecules could guide EPIC-T cell trafficking *in vivo*.

In summary, we demonstrate that primary human T cells can produce functional VSV-G pseudo-typed Cas9 EDVs, and we identified and resolved bottlenecks in their production, including the optimization of primary T cell engineering and Gag architectures. We further validated a non-viral site-specific knock-in approach to generate EPIC-T cells. We anticipate that cell-based delivery modalities will serve as valuable additions to existing delivery methods of gene editors and aid in translation of gene editing therapies.

## Limitations of the Study

We anticipate that the efficiency of EPIC-T cell production could be further improved in the future. We anticipate that improvements in large sequence integration with serine integrases^37^, bridge recombinases^38^, quad prime editing^39^, and other approaches will aid in the ability to knock-in EDV production machinery. Additionally, retroviral Gag can be immunogenic, and further investigation of *in vivo* persistence of EPIC-T cells in immune-competent animals will be required. We hypothesize that, if EDV machinery proves immunogenic, that alternative non-Gag based exosome loading systems, trogocytosis approaches, inducible expression, or masking approaches used for allogeneic cell therapies^40,41^ may mitigate the risk of immunologic rejection.

## Supplemental Figures

**Figure S1.**
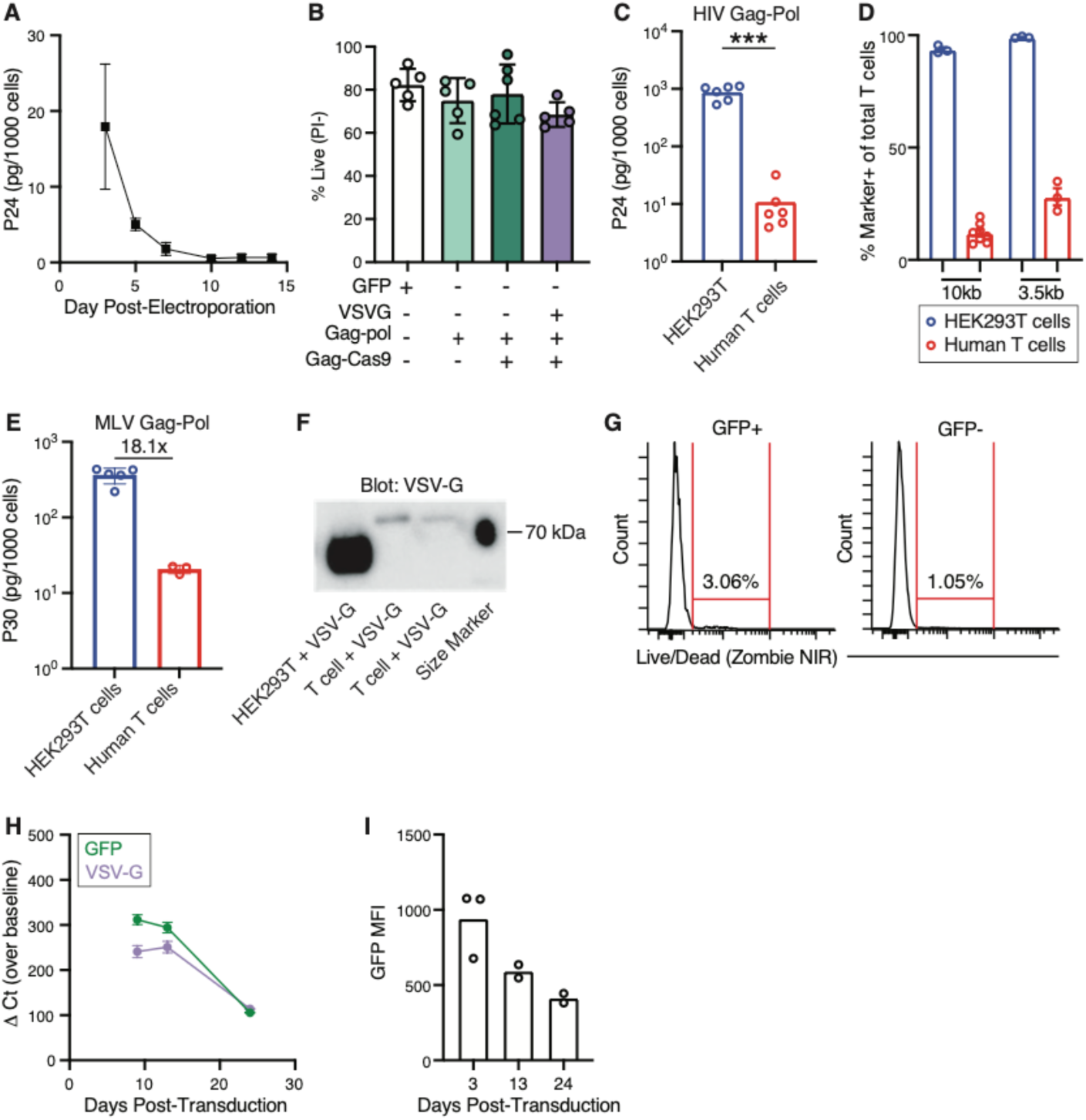
Primary human immune cells can produce a variety of enveloped delivery vehicles (EDVs). **(A)** Levels of HIV P24 detected over time in the supernatant of primary human T cells nucleofected with plasmid encoding HIV Gag-Pol, normalized to cell numbers. **(B)** Percent of live primary human T cells nucleofected with plasmids encoding the indicated genes as measured by propidium iodide staining by flow cytometry on day 3 post nucleofection. **(C)** Levels of HIV P24 detected in the supernatant of HEK293T cells or primary human T cells following introduction of plasmid encoding HIV Gag-Pol, normalized to cell numbers. P-values from two-sided Student’s t-test. **(D)** Percent of HEK293T cells (blue) or primary human T cells (red) expressing fluorescent markers encoded by a 10 kb plasmid (mCherry) or a 3.5 kb plasmid (GFP). **(E)** Levels of MLV P30 detected in the supernatant of HEK293T cells or primary human T cells nucleofected with plasmid encoding MLV Gag-Pol, normalized to cell numbers. **(F)** Levels of VSV-G protein expressed by HEK293T cells or two biological replicates of primary human T cells on day 10 after lentiviral transduction of GFP-IRES-VSV-G measured by Western blot. **(G)** Percent of dead GFP+ and GFP-primary human T cells after lentiviral transduction of GFP-IRES-VSV-G as measured by flow cytometry. **(H)** Levels of GFP and VSV-G transcript expressed by primary human T cells over time after lentiviral transduction of GFP-IRES-VSV-G measured by qPCR. **(I)** GFP median fluorescence intensity in primary human T cells over time after lentiviral transduction of GFP-IRES-VSV-G measured by flow cytometry. Abbreviations: HIV = Human immunodeficiency virus. Gag-Pol = Group-specific antigen and polymerase polyprotein. MLV = Moloney murine leukemia virus. VSV-G = Vesicular Stomatitis Virus Glycoprotein. GFP = Green fluorescent protein. qPCR = quantitative polymerase chain reaction. *** denotes nominal P-value < 0.005.

**Figure S2.**
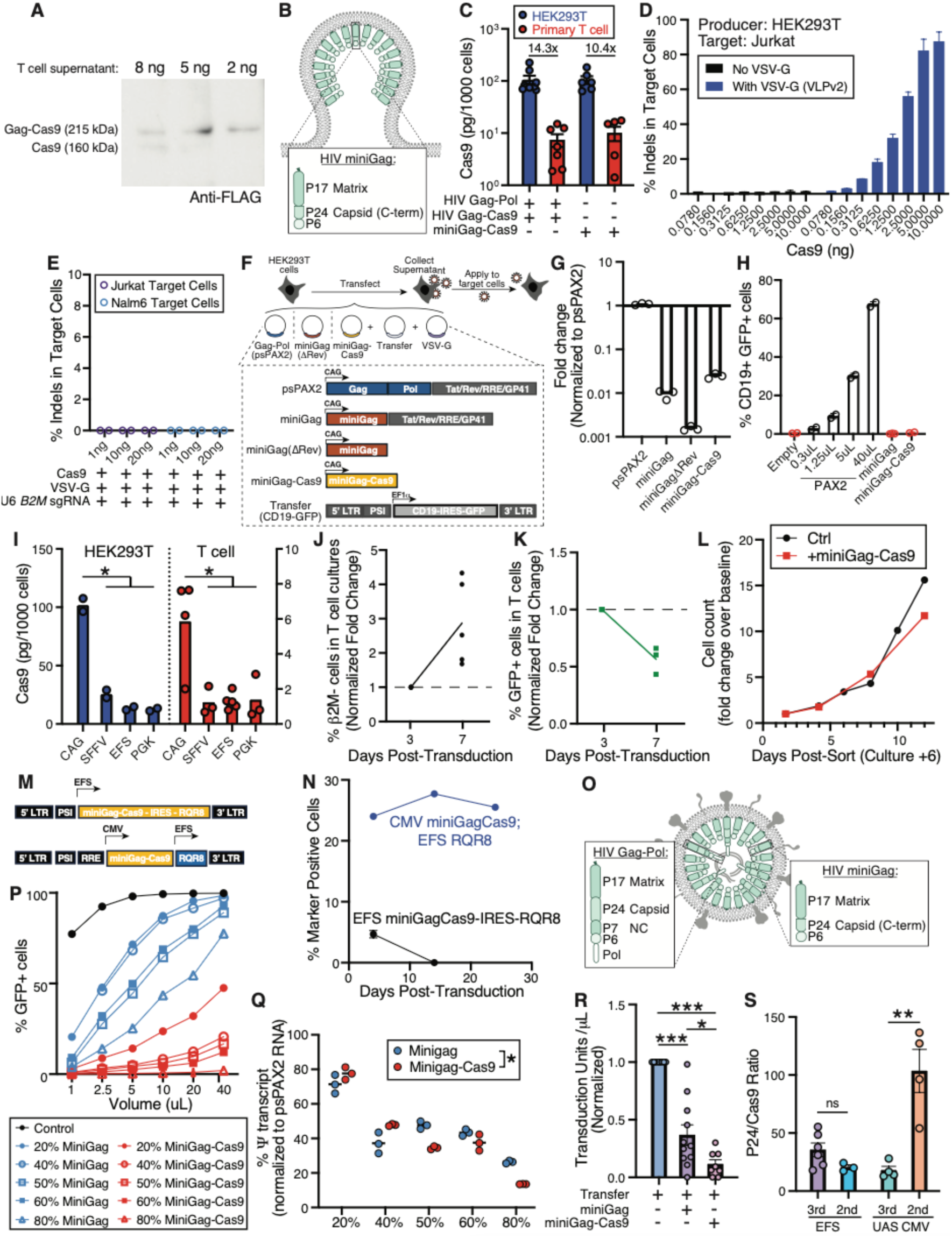
T cells can produce EDVs that package functional Cas9 ribonucleoproteins capable of gene editing in target cells. **(A)** Levels of FLAG-tagged Gag-Cas9 fusion protein and Cas9 protein in supernatants of primary human T cells after nucleofection with plasmids encoding HIV Gag-Pol and HIV Gag-Cas9 measured by Western blot. Nanogram amounts denote detected levels of P24 in loaded sample. **(B)** Schematic of HIV-based miniGag on an EDV. **(C)** Levels of Cas9 protein detected in the supernatant of HEK293T cells transfected with, or primary human T cells nucleofected with, plasmids encoding the indicated genes, normalized to cell numbers. **(D)** Percent of *B2M* gene amplicons with indels in Jurkat target cells incubated with supernatants from HEK293T cells transfected with plasmids encoding HIV Gag-Pol and Gag-Cas9, with or without plasmid encoding VSV-G. Cas9 levels in the supernatant were quantified by ELISA and used to calculate the quantity of supernatant transferred as indicated. **(E)** Percent of *B2M* gene amplicons with indels in Jurkat or Nalm6 target cells incubated with supernatants from primary human T cells nucleofected with plasmids encoding alone, VSV-G, and *B2M* sgRNA. Cas9 levels in the supernatant were quantified by ELISA and used to calculate the quantity of supernatant transferred as indicated. **(F)** Schematic of transfection of HEK293T cells with plasmids encoding HIV Gag-Pol, miniGag, or miniGag-Cas9, together with transfer plasmid and VSV-G. Supernatants were collected and transferred onto HEK293T target cells. **(G)** CD19 transcript as measured by qPCR in the supernatant of HEK293T cells transfected with the indicated genes with CD19-GFP transfer plasmid and VSV-G, normalized to levels of cells transduced with second generation lentiviral packaging vector (psPAX2). **(H)** Percent of HEK293T target cells expressing CD19 and GFP (encoded on the lentiviral transfer plasmid) after incubation with the indicated volume of supernatant from HEK293T packaging cells transfected with plasmids encoding the indicated genes with CD19-GFP transfer plasmid and VSV-G, as measured by flow cytometry. For Empty, miniGag, and miniGag-Cas9 conditions, 100µL supernatant was added. **(I)** Levels of Cas9 protein detected in the supernatant of HEK293T cells (left) or primary human T cells (right) after nucleofection with plasmid encoding the miniGag-Cas9 expressed under the control of the indicated promoter. **(J)** Flow cytometry detection of % β2M*-*negative cells (surface expressed), normalized for fold change over baseline, among primary human T cells over time after transduction with lentivirus encoding miniGag-Cas9 and lentivirus encoding *B2M* sgRNA, VSV-G, and GFP. **(K)** Flow cytometry detection of % GFP-positive cells, normalized for fold change over baseline, among primary human T cells over time after transduction with lentivirus encoding miniGag-Cas9 and lentivirus encoding *B2M* sgRNA, VSV-G, and GFP. **(L)** Fold-change over time in numbers of sorted primary human T cells following transduction with GFP control virus or lentivirus encoding miniGag-Cas9 (denoted in following schematic). **(M)** Schematic of bicistronic transfer cassette encoding miniGag-Cas9 and RQR8 under control of the EFS promoter and dual-promoter transfer cassette encoding miniGag-Cas9 under control of the CMV promoter and RQR8 under control of the EFS promoter. **(N)** Percent of primary human T cells expressing the RQR8 marker over time after transduction with each lentivirus shown in (L), as measured by flow cytometry. **(O)** Schematic of mosaic lentivirus produced by a combination of HIV Gag-Pol and HIV miniGag-Cas9. **(P)** Percent of target HEK293T cells expressing GFP after incubation with supernatant from HEK293T packaging cells co-transfected with varying ratios of psPAX2 packaging plasmid and miniGag or miniGag-Cas9, with VSV-G and a third-generation lentiviral transfer plasmid encoding CD19 and GFP. **(Q)** Packaging of viral RNA as detected by qPCR of the Psi packaging sequence in supernatant of HEK293T packaging cells following transfection of varying ratios of psPAX2 packaging plasmid and miniGag or miniGag-Cas9, normalized to levels of cells transfected with psPAX2 packaging plasmid alone. P-value determined by 2-way ANOVA. **(R)** Transduction units per μL of lentiviruses produced by HEK293T cells transfected with transfer plasmids encoding miniGag or miniGag-Cas9, normalized to levels from HEK293T cells transfected with GFP control transfer plasmid. **(S)** Ratio of P24 HIV capsid protein to Cas9 protein in lentivirus produced by HEK293T cells transfected with 2^nd^ or 3^rd^ generation transfer plasmids encoding miniGag-Cas9 under the control of the EFS promoter (constitutive) or a UAS-minimal CMV promoter (inducible). Abbreviations: HIV = Human immunodeficiency virus. Gag-Pol = Group-specific antigen and polymerase polyprotein. VSV-G = Vesicular Stomatitis Virus Glycoprotein. *B2M* = Beta-2-microglobulin gene. sgRNA = single guide ribonucleic acid. indel = insertion or deletion mutation. GFP = Green fluorescent protein. qPCR = quantitative polymerase chain reaction. IRES = internal ribosome entry site. EFS = Elongation Factor 1 alpha short. UAS = upstream activation sequence. CMV = cytomegalovirus. * denotes nominal P-value < 0.05. ** denotes nominal P-value < 0.01. *** denotes nominal P-value < 0.005.

**Figure S3.**
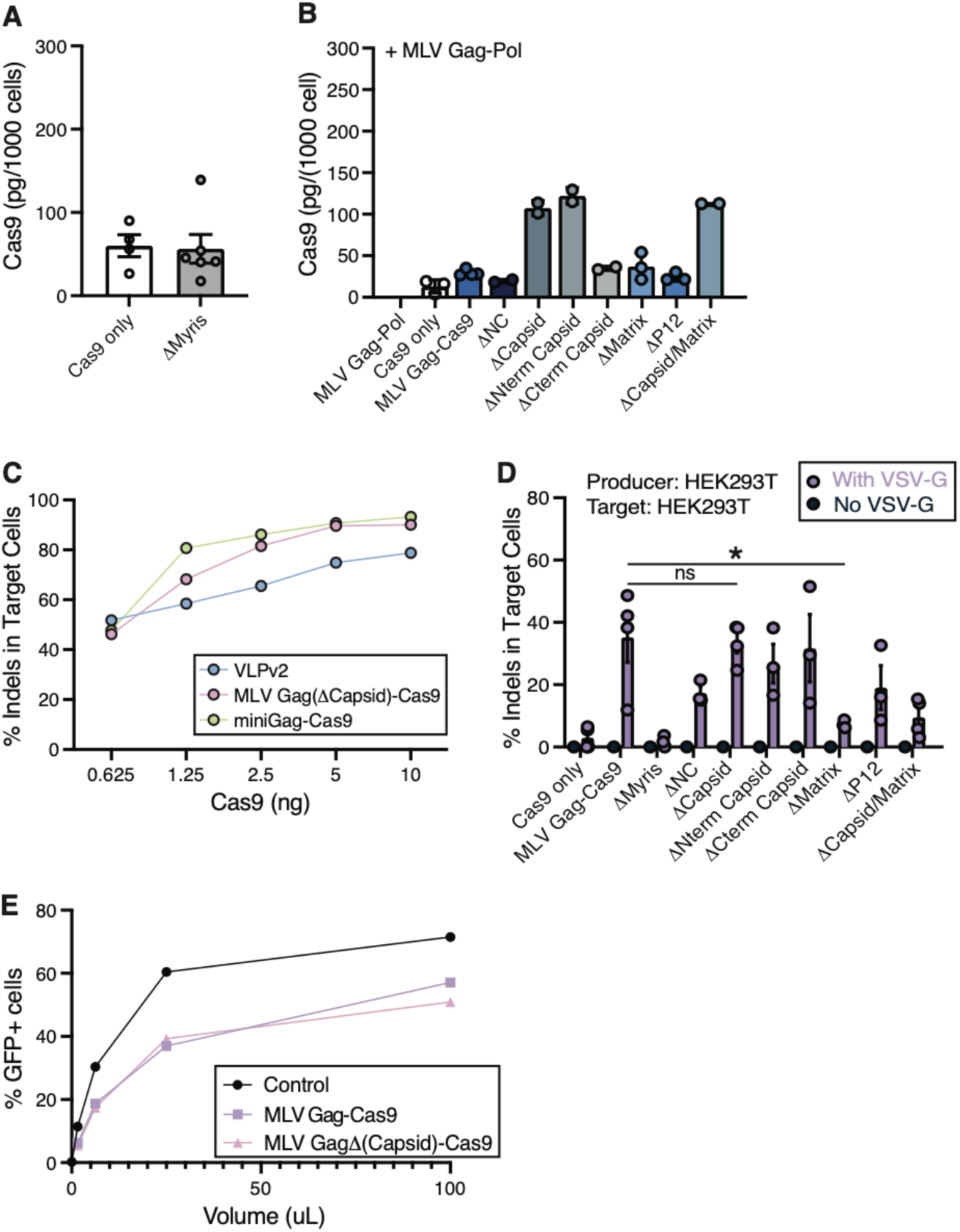
Development of a miniaturized MLV-based EDV system produced efficiently by EPIC-T cells. **(A)** Levels of Cas9 protein detected in the supernatant of HEK293T cells transfected with plasmid encoding Cas9 only or MLV Gag(ΔMyris)-Cas9. **(B)** Levels of Cas9 protein detected in the supernatant of HEK293T cells transfected with plasmid encoding MLV Gag-Pol together with plasmid encoding the indicated gene, normalized to cell numbers. Δ indicates a deletion of the indicated region from MLV Gag-Cas9 fusion gene. **(C)** Percent of *B2M* gene amplicons with indels in Jurkat target cells incubated with varying amounts of supernatants from HEK293T cells transfected with plasmids encoding HIV Gag-Pol, Gag-Cas9 and VSV-G (VLPv2), or transfected with MLV Gag(ΔCapsid)-Cas9 or miniGag-Cas9 together with VSV-G and sgRNA. Cas9 levels in the supernatant were quantified by ELISA and used to calculate the quantity of supernatant transferred as indicated. **(D)** Percent of *B2M* gene amplicons with indels in HEK293T target cells incubated with supernatants from HEK293T cells transfected with plasmids encoding the indicated genes with plasmid encoding sgRNA and with or without plasmid encoding VSV-G. **(E)** Percent of HEK293T cells expressing GFP after incubation with supernatant from HEK293T packaging cells co-transfected with psPAX2 packaging plasmid encoding MLV Gag-Cas9, MLV Gag(ΔCapsid)-Cas9, or control with VSV-G and a third-generation lentiviral transfer plasmid encoding CD19 and GFP. Abbreviations: MLV = Moloney murine leukemia virus. Gag-Pol = Group-specific antigen and polymerase polyprotein. *B2M* = Beta-2-microglobulin gene. HIV = Human immunodeficiency virus. VSV-G = Vesicular Stomatitis Virus Glycoprotein. indel = insertion or deletion mutation. GFP = Green fluorescent protein. * denotes nominal P-value < 0.05.

**Figure S4.**
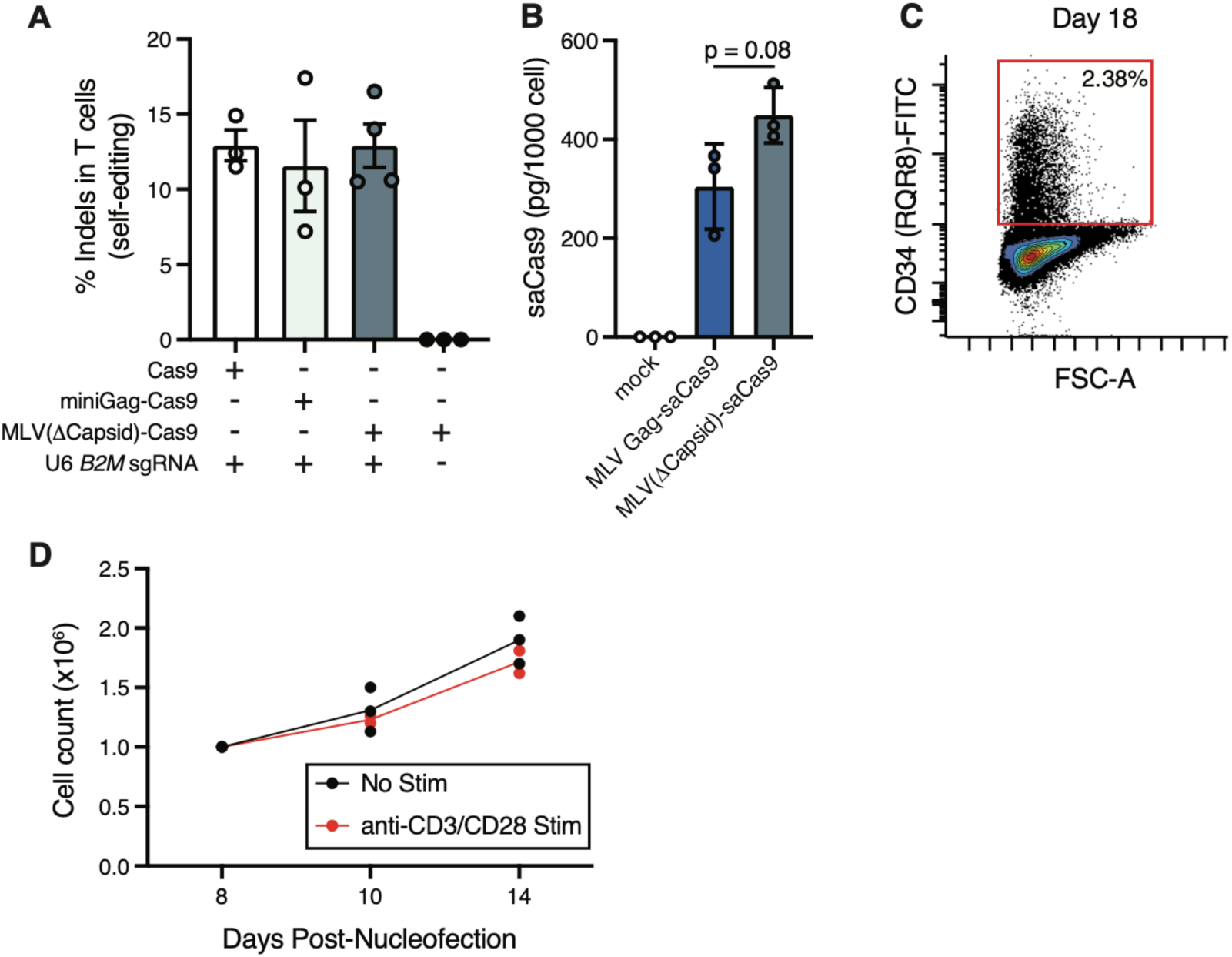
Generation of EPIC T cells via all-in-one, site-specific knock-in of miniaturized MLV EDV production machinery. **(A)** Percent of *B2M* gene amplicons with indels in EPIC-T cell cultures generated by nucleofection of the indicated genes. **(B)** Levels of saCas9 protein detected in the supernatant of primary human T cells nucleofected with plasmids encoding the indicated genes, normalized to cell numbers. **(C)** Expression of the RQR8 marker encoded in the HDRT of EPIC-T cells produced by non-viral knock-in after 18 days in culture. **(D)** Number of EPIC-T cells manufactured by site-specific knock-in with and without anti-CD3/CD28 stimulation over time. Abbreviations: MLV = Moloney murine leukemia virus. *B2M* = Beta-2-microglobulin gene. indel = insertion or deletion mutation. saCas9 = *Staphylococcus aureus* Cas9. HDRT = homology-directed repair template. EPIC-T cell = Enveloped Particle Information Courier-T cell.

## Methods

### Plasmid generation

VLPv2 generation plasmids JRH-1345 U6-B2M sgRNA Gag-Cas9 v2 (Addgene plasmid #201916) and pJRH-1346 U6-B2M sgRNA Gag-pol v2 (Addgene plasmid # 201917) and MiniEDV generation plasmid pWN_U6-B2M-miniGag-Cas9 (Addgene plasmid #228958) were gifts from Jennifer Doudna. MLV-based eVLP generation plasmids v5 eVLP gag-pro-pol (Addgene plasmid #228466) and pCMV-MMLVgag-3xNES-Cas9 (Addgene plasmid #181752) were gifts from David Liu. ENVLPE+ EDV generation plasmids pCMV_ENVLPE+(Addgene plasmid #232427), pU6_B2M_sgRNA(PP7) (Addgene plasmid # 232438), and pCMV_miniENVLPE (Addgene plasmid #23243) were gifts from Gil Westmeyer. PsPAX2 (Addgene plasmid #12260) and pMD2.G (Addgene plasmid #12259) were gifts from Didier Trono. Transfer plasmids for EFS miniGag-Cas9 and U6 *B2M* sgRNA, EFS VSV-G were custom ordered from VectorBuilder using the annotated sequences from Addgene plasmid # 228958 and pMD2.G. For supplemental figure 1, the 3.5kb plasmid was pmaxGFP obtained from Lonza. For Supplemental Figure 2, the transfer plasmid used for mosaic experiments was pJRH-1363 LV EF1a-CD19 IRES-EGFP (Addgene plasmid #201919)

Gibson cloning was used to produce all remaining plasmids. We introduced the *B2M* sgRNA targeting sequence *gagtagcgcgagcacagcta* from Addgene plasmid #201916 into Addgene plasmid #232438 and cloned the cassette upstream of spCas9 in the pCAGGS expression plasmid. The 10kb plasmid (Figure S1B) was produced by cloning an mCherry reporter into Addgene plasmid #228958. The GFP-IRES-VSVG plasmid (Figure 1 and Supplemental Figure 1) was generated by cloning GFP and VSVG into a 3^rd^ generation lentivirus transfer backbone. GFP-IRES-MiniGag and GFP-IRES-Cas9 (Figure 2) were generated by cloning miniGag and Cas9 from Addgene plasmid #228958 into a 3^rd^ generation lentiviral transfer plasmid containing GFP–IRES. The anti-silencing EFS, HBBIVS2, minigag-Cas9 plasmid was cloned by placing the HBBIVS2 intron from pSA128 A2UCOE-HBB_IVS2-Cas9-BFP Addgene plasmid #249152; gift from Michael Bassik) in the 5’ UTR of the transfer plasmid containing EFS – miniGag-Cas9. For Supplemental Figure 2, the U6 – *B2M* sgRNA, Cas9 expression plasmid was generated by Gibson deletion of miniGag and linker from Addgene plasmid #228958. The promoters tested in Supplemental Figure 2 were ordered as gBlocks from IDT using published consensus sequences and cloned in place of the CAG promoter present in Addgene plasmid #228958. UAS_mincmv_ was obtained from pHR-Gal4UAS-PGK-BFP (Addgene plasmid #247505, a gift from Wendell Lim) and cloned into the miniGag-Cas9 transfer plasmid where EFS was placed in front of RQR8. The 2nd generation lentiviral transfer plasmid used was the pHR backbone from Addgene plasmid #247505 and the referenced UAS_mincmv_ was utilized in place of EFS where indicated. MiniGagΔRev was cloned using Gibson deletion of intervening segments from Addgene plasmid #228958. All MLV Gag-Cas9 mutants were generated from Addgene plasmid #181752 by Gibson deletion of the indicated Gag subunit. For MLV Gag-Cas9 lentiviral transfer, we cloned the desired sequence from Addgene plasmid # 181752 and MLV GagΔCapsid mutant into a 3^rd^ generation transfer plasmid. For staphylococcus aureus Cas9, we obtained the sequences from pX601-AAV-CMV::NLS-SaCas9-NLS-3xHA-bGHpA;U6::BsaI-sgRNA (Addgene plasmid # 61591, gift from Feng Zhang) and cloned into Addgene plasmid # 81752 and in our MLV GagΔCapsid mutant in in place of spCas9. NEB STBL3 was transformed by all clones. Plasmids were prepared with the ZymoPURE II Plasmid Midiprep Kit. All plasmids were sequence confirmed by Azenta before use.

### Tissue culture

Lenti-X cells were purchased from Takara and cultured in DMEM (Corning) supplemented with 10% fetal bovine serum (Omega scientific), 1% penicillin-streptomycin (Gibco), and Gluta-Max (Gibco) (cDME). Jurkat cells were obtained and authenticated by the UC San Francisco Cell Culture Facility and cultured in RPMI 1640 (Corning) with 10% fetal bovine serum (Omega Scientific), 1% penicillin-streptomycin (P/S, Gibco), and Gluta-Max (Gibco) (cRPMI). A549 and Nalm6 cells were a gift from J. Eyquem and cultured in Ham’s F12K (ATCC) supplemented with 10% fetal bovine serum (Omega scientific), 1% penicillin-streptomycin (Gibco), and Gluta-Max (Gibco) and cRPMI respectively. A204 was a gift from N. Cho and cultured in McCoy’s 5A (Gibco) supplemented with 10% fetal bovine serum (Omega scientific), 1% penicillin-streptomycin (Gibco) and Gluta-Max (Gibco). All cell lines were maintained in a 37°C, 5% CO_2_ incubator and checked for mycoplasma contamination by ATCC Universal Mycoplasma Detection Kit (30-1012K)

### Primary T cell isolation and culture

Human peripheral blood mononuclear cells (PBMCs) were isolated from deidentified healthy platelet pheresis leukoreduction filters (Vitalant) using density centrifugation and cryopreserved until use. CD3, CD4, and CD8 T cells were isolated from PBMCs by negative selection using EasySep Human T Cell Isolation Kit (StemCell, 17951), EasySep Human CD4+ T cell Isolation Kit (StemCell, 17952), and EasySep Human CD8+ T cell Isolation Kit (StemCell, 17953) per manufacturer’s instructions. Isolated T cells were cultured at 1*10^6^ cells per mL in complete RPMI (RPMI 1640 (Corning) with 10% fetal bovine serum (Omega Scientific), 1% P/S, NEAA, 25mM HEPES, Sodium pyruvate, and Glutamax (Gibco)) supplemented with 100 U/mL IL-2 (Peprotech), 5 ng/mL IL-7 (Peprotech) and 5 ng/mL IL-15 (Peprotech) (T cell media) and rested either one hour or overnight in a 37°C, 5% CO_2_ incubator in before 1:1 stimulation with Dynabeads Human T-Activator CD3/CD28 (Gibco). T cells were stimulated for 48-72 hours before subsequent downstream nucleofection or lentiviral transduction. Unless otherwise noted in figure legend, T cells were split were 2-3 days to maintain a 1*10^6^ cells/mL culture density.

### Primary Human Monocyte Culture

Monocytes were isolated from PBMCs using EasySep Human Monocyte Isolation Kit (StemCell, 19359) and cultured at 1*10^6^ cells/mL in complete RPMI supplemented with 20ng/mL GM-CSF (Peprotech) and IL-4 (Peprotech). Monocytes were maintained at 1*10^6^ cells/mL density and cultured for one week before nucleofection.

### Plasmid Nucleofection

48-72 hours post-stimulation, T cells were separated from Dynabeads with an EasySep Magnet (StemCell). Cells were counted on a Countess 2 FL Automated Cell Counter (Thermo Fisher) using Trypan Blue dye to determine viability. 1.5*10^6^ T cells were centrifuged at 200 × g for 8 minutes and resuspended in 20 µL P3 Buffer (Lonza, V4SP-3960) before being mixed with prepared plasmid. Nucleofection was performed on a Lonza 4D Nucleofector Core unit (Lonza, AAF-1003B) paired with a 96-well Unit (Lonza, AAF-1003S) using pulse code EH-115 for T cells, DP-148 (P3 Buffer) for monocytes, and CL-120 (Buffer SE) for Jurkat cells. Following pulse, 100 µL of prewarmed culture media was added, and cells were allowed to recover on cuvette plate for 15 minutes before transfer to culture plates at 1*10^6^ cells/mL density. 72 hours following nucleofection, supernatant was collected for analysis. For VLPv2 generation, 1.25µg of HIV Gag-Pol, 1.25µg Gag-Cas9, 500ng pMD2.G was utilized. For miniEDVs, 1.25 µg miniGag-Cas9 and 500ng pMD2.G was added. For MLV Gag-Cas9 and mutant experiments, 1.25 µg of Gag-Cas9 construct was used with 500ng pMD2.G with and without 1.25 µg MLV Gag-Pol. For ENVLPE+, 1.25 µg of ENVLPE+ or miniENVLPE was co-nucleofected with 1.25 µg U6 sgRNA PP7, Cas9, and 500ng pMD2.G if indicated.

### Lentivirus generation

1*10^7^ Lenti-X cells were seeded in a 10cm plate the afternoon before transfection. In the morning, each 10cm plate was checked for 80-90% confluency before proceeding. 50μL lipofectamine 3000 (Thermo Fisher) in 1mL Opti-MEM was added to 1mL DNA mixtures consisting of 10μg psPAX2, 10μg transfer plasmid, 4μg pMD2.G, and 48μL P3000 in Opti-MEM (Gibco). These values were adjusted proportionally based on surface area if virus was generated with a different culture vessel (6 well plate for pilot, T-175 flask for larger preparations). For all transfer plasmids containing VSV-G including *GFP-IRES-VSVG* and *U6 B2M sgRNA, VSV-G,* 15μg of third generation packaging system (LVMAX, Thermo Fisher A43237) was used per 10μg of transfer plasmid. 6 hours after transfection, media was exchanged for fresh cDMEM supplemented with ViralBoost (Alstem). Supernatant was collected 48 and 72 hours post transfection, spin clarified 500g for 5 minutes, filtered through a 0.45μm PES membrane, and mixed with Lenti-X concentrator (Takara) according to manufacturer’s instructions overnight, and concentrated 100x initial volume in Opti-MEM and either used fresh or aliquoted and stored at -80°C. For functional titer calculation, we added 10μL of 100x concentrated viral preparations to 1*10^6^ stimulated primary human T cells in 1mL T cell culture media, removed virus-containing media after 24 hours, and assayed to assess % marker positive cells 96 hours after initial virus application to determine transducing units per virus volume.

For the mosaic virus tests with miniGag-Cas9, miniGag, MLV Gag, and MLV GagΔCapsid, we seeded 1.8*10^6^ Lenti-X cells in 6 well plates and transfected with 625ng psPAX2, 1.25μg CD19-GFP transfer plasmid, 500ng pMD2.G, and varying amounts of the mosaic gag construct (156.25ng miniGag-Cas9 for 20% mosaicism, 417 ng for 40%, 625 ng for 50%, 937.5ng for 60%, 2.5 μg for 80%) with 2μL/μg of DNA of lipofectamine 3000 and P3000. 6 hours after transfection, media was replaced with cDMEM, and mosaic virus preparations were collected 48 hours after transfection and applied to 25,000 Jurkat cells in 100μL cRPMI in a 96 well u-bottom plate.

### ELISA

EDV and control supernatant was spun 500g x 5 minutes and filtered through a 0.45μm PES membrane. For p24 analysis of primary cell samples, 100μL of unconcentrated, undiluted supernatant was analyzed by Lentivirus-associated p24 ELISA Kit (Cell Biolabs) according to manufacturer’s instructions. For p30 and Cas9 T cell supernatant ELISA, 180μL of unconcentrated, undiluted supernatant was digested with 20μL of Triton-X solution (Cell Biolabs) for 30 minutes at 37°C before analysis with MuLV Core Antigen ELISA Kit or Cas9 (CRISPR Associated Protein 9) ELISA Kit (S. pyogenes) (Cell Biolabs) per manufacturer’s instructions in technical duplicate. Occasionally, ELISA samples were aliquoted and frozen prior to thaw and Triton-X digestion. For p24 and p30 quantification, HEK293T and Jurkat samples were diluted 1:1000 before analysis. For Cas9 quantification, HEK293T samples were diluted 1:10. For all ELISA kits, absorbance at 450 nm was measured by a plate reader (Biotek). P24, p30, and Cas9 amounts were calculated by comparison to a serial dilution of the relevant standard. The total protein amount was calculated by dividing by the sample volume run for the assay (90μL for Cas9) and multiplying by the total supernatant volume harvested (variable). This total was then normalized based on the cell count at time of harvest.

### HEK293T Cas9-EDV production

All Cas9 EDVs were generated by seeding 1.8*10^6^ Lenti-X cells in 6 well plates and transfecting the next day with 2μL/μg lipofectamine 3000 and P3000. For VLPv2, we transfected 1.25μg JRH-1345, 1.25μg JRH-1346, and 500ng pMD2.g. For miniEDV, 1.25μg WN miniGag-Cas9 plus and 500ng pMD2.g. For MLV full length and mutant EDVs, 1.25μg MLV Gag-Cas9 (and all individual mutants) plus or minus 1.25μg MLV Gag-Pol, 1.25μg of U6 *B2M* sgRNA plasmid, and 500ng pMD2.G were transfected. 6 hours after transfection, media was exchanged for fresh cDMEM, and Cas9 EDV supernatants were collected 48 and 72 hours post transfection. Collected supernatant was spun 500g x 5 minutes, filtered through a 0.45μM PES filter, aliquoted and either used fresh or stored at -80°C.

### Quantitative RT–PCR

1*10^6^ T cells were pelleted, lysed with buffer RLT, and RNA was extracted with Qiagen RNeasy Kit according to the manufacturer’s instructions. Quantitative RT-PCR was performed using PrimeTime One-Step RT–qPCR Master Mix (IDT) and custom qPCR primers and probe ordered from IDT to detect GFP or VSVG. Reaction was run on a Bio-Rad C1000 Touch Thermocycler with CFX384 Optical Reaction Module. For Psi sequence detection, mosaic lentivirus was harvested and concentrated as previously described and 100μL of concentrated sample was added to Buffer RLT, and RNA was extracted with Qiagen RNeasy Kit. Custom qPCR primers and probe to detect Psi were ordered from IDT.

### Flow Cytometry

For GFP expression (Figure 1L; S1B, G, I; 2M; S2G, J, N; S3G), cells were stained with Zombie NIR (1:200) in PBS containing 1% bovine serum albumin. For data shown in Figure S1D, cells were stained with Propidium Iodide (1:20). For CD19 expression (Figure S2G), cells were stained with anti-human CD19 BV605 (1:100). For Figure 2D, cells were stained with anti-human B2M-FITC (1:100). For Figures 2M and 4C, cells were stained 1:5 with CD34-FITC. All stained cells were analyzed on a LSR Fortessa Cell Analyzer with 96-well autosampler (BD).

### Western blot analysis

For anti-Flag detection of Gag-Cas9 and Cas9, T cell supernatant was mixed with Laemmli buffer containing 10% molecular biology grade 2-mercaptoethanol and heated at 90°C for 5 min. For anti-VSV-G detection, pelleted T cells were lysed with cold RIPA + anti-protease cocktail (Cell Biolabs) on ice for 15 minutes. The mixture was centrifuged at max speed for 10 min at 4°C, and supernatant was transferred to a new tube, and protein amount was quantified by Bradford Assay. Samples were resolved on a 4%–20% SDS–PAGE gel (Invitrogen) and transferred (20V for 16 hours at 4°C) to a PVDF membrane. Membranes were blocked with 5% milk for 2 hours at room temperature and incubated with primary antibody at 4°C overnight following by secondary antibody at room temperature for 1 hour. SuperSignal West Pico Plus (Thermo Fisher) was added for 5 minutes. Membranes were washed, and imaging was performed using a BioRad Chemidoc.

### Amplicon sequencing

Target cells were plated at 2.5*10^4^ in 100μL in a 96 well u-bottom plate with 4μg/mL protamine sulfate. The next day, producer generated EDV supernatants at known Cas9 amounts, not exceeding 100μL volume, were added. 24 hours after EDV application, the media was changed. 72 hours after EDV application, media was removed, cells were washed once with PBS, and genomic DNA was extracted by resuspending in 50μL QuickExtract (Lucigen), transferring to PCR tubes, and heating on thermocycler at 65°C for 20 minutes and 95°C for 20 minutes. Q5 Hi Fidelity polymerase (NEB) was used to amplify the Cas9 *B2M* sgRNA target site using 5’ TCACCCAGTCTAGTGCATGC and 5’ GACGCTTATCGACGCCCTAA. PCR products were resolved on a 1.5% agarose gel, and fragments were purified using GeneJET PCR Purification Kit. PCR products were submitted to Azenta for sanger sequencing. Trace files were visualized on TIDE: Tracking of Indels by Decomposition and % indels were quantified by Synthego ICE.

### Site Specific Knock-in

Custom sgRNA was resuspended at 80 µM in IDT duplex buffer and stored in single-use aliquots at -80°C. Cas9 mRNA (1mg/mL) was ordered from Synthego. Nanoplasmid HDRT template was ordered from Aldevron. 1 µg of Cas9 mRNA, 1 µg of sgRNA, and 2 µL of plasmid DNA template at 1 ug/µL were mixed in a 96 well u-bottom plate with 20 µL activated T cells (1.5*10^6^ total), transferred to cuvette, and nucleofected with Lonza 4D Nucleofector Core Unit (Lonza, AAF-1003B) paired with a 96-well Unit (Lonza, AAF-1003S) using pulse code EH-115. After recovery on plate, cells were plated in desired culture vessel with T cell Media with 5 μM M3814 (MedChemExpress) DNA PK inhibitor and 5 μM ART558 (MedChemExpress) Polθ inhibitor. Inhibitor-containing media was removed after 24 hours. Nucleofected T cells were split every 2-3 days to maintain 1*10^6^ cells/mL density.

### Statistical analysis

Statistical analysis was performed using Prism v.10. Reported P-values are nominal. Details can be found in the figure legends corresponding to each figure panel.

## Data availability

Sequencing and flow cytometry raw data files are available upon request. All other data are available in the main text or the Supplementary Information.

## Acknowledgements

We thank all members of the Spitzer laboratory, J. Eyquem, A. Marson, S. Dodgson for their thoughtful input on this project. Funding was provided to James Meixiong by 5T32AR007175-48 and by the following awards to M.H.S.: American Association for Cancer Research award 20-20-01-SPIT, Cancer Research Institute award CRI4437, NIH awards R01DE032033 and R01CA290027, American Cancer Society award RSG-22-141-01-IBCD, DOD US Army Med. Res. Acq. Activity Award BC220499, CRISPR Cures for Cancer, and the Weill Cancer Hub West, as well as UCSF Diabetes Research Center grant NIH P30DK135103.

## Author contributions

J. M. and M.H.S. conceptualized this work. J. M., L.C., and B.S. developed the methodology. J. M. and O.Z. carried out the investigation. J. M. and M.H.S. created visual representations, wrote the initial draft, edited the manuscript. M.H.S. was responsible for funding acquisition and project administration.

